# Benchmark of data processing methods and machine learning models for gut microbiome-based diagnosis of inflammatory bowel disease

**DOI:** 10.1101/2021.05.03.442488

**Authors:** Ryszard Kubinski, Jean-Yves Djamen-Kepaou, Timur Zhanabaev, Alex Hernandez-Garcia, Stefan Bauer, Falk Hildebrand, Tamas Korcsmaros, Sani Karam, Prévost Jantchou, Kamran Kafi, Ryan D. Martin

**Author notes:** Co-corresponding authors - send correspondence to or.

## Abstract

**Background:** Inflammatory bowel disease (IBD) patients wait months and undergo numerous invasive procedures between the initial appearance of symptoms and receiving a diagnosis. In order to reduce time until diagnosis and improve patient wellbeing, machine learning algorithms capable of diagnosing IBD from the gut microbiome’s composition are currently being explored. To date, these models have had limited clinical application due to decreased performance when applied to a new cohort of patient samples. Various methods have been developed to analyze microbiome data which may improve the generalizability of machine learning IBD diagnostic tests. With an abundance of methods, there is a need to benchmark the performance and generalizability of various machine learning pipelines (from data processing to training a machine learning model) for microbiome-based IBD diagnostic tools.

**Results:** We collected fifteen 16S rRNA microbiome datasets (7707 samples) from North America to benchmark combinations of gut microbiome features, data normalization methods, batch effect reduction methods, and machine learning models. Pipeline generalizability to new cohorts of patients was evaluated with four binary classification metrics following leave-one dataset-out cross validation, where all samples from one study were left out of the training set and tested upon. We demonstrate that taxonomic features obtained from QIIME2 lead to better classification of samples from IBD patients than inferred functional features obtained from PICRUSt2. In addition, machine learning models that identify non-linear decision boundaries between labels are more generalizable than those that are linearly constrained. Prior to training a non-linear machine learning model on taxonomic features, it is important to apply a compositional normalization method and remove batch effects with the naive zero-centering method. Lastly, we illustrate the importance of generating a curated training dataset to ensure similar performance across patient demographics.

**Conclusions:** These findings will help improve the generalizability of machine learning models as we move towards non-invasive diagnostic and disease management tools for patients with IBD.

## Introduction

The human gut microbiome is a collection of microbes, viruses, and fungi residing throughout the digestive tract. The gut microbiota plays an important role in human health, influencing food digestion, the immune system, mental health, and numerous other functions (reviewed in [1]). In line with the functional role in human health, alterations in the gut microbiome have been linked to illnesses such as multiple sclerosis, type II diabetes, and inflammatory bowel disease (IBD) [2, 3]. IBD comprises two main subtypes: Crohn’s disease (CD) and ulcerative colitis (UC), characterized by periodic inflammation throughout the gastrointestinal tract or localized to the colon, respectively [4]. The prevalence of IBD is increasing globally over the last several decades, from 79.5 to 84.3 per 100 000 people between 1990 and 2017, with Canada having among the highest IBD rates at 700 per 100 000 people in 2018 [5, 6]. Although the disease etiology is currently undetermined, the increasing rates of IBD have been linked to lifestyle factors, such as a Western diet [7].

Currently, IBD diagnosis and monitoring is primarily performed via blood tests, fecal calprotectin, and endoscopies. These methods can be costly, invasive, and display variable accuracy, all of which leads to delayed diagnosis and infrequent disease monitoring [8]. Therefore, there is an unmet need for the development of further non-invasive, low-cost, and rapid methods for screening, diagnosis, and disease management for the growing number of IBD patients [9, 10]. One potential diagnostic test within these constraints involves using the gut microbiome composition to identify patients with IBD.

Over the past decade, several studies have compared the gut microbiome profiles of healthy individuals and those with CD or UC [2, 11–21]. Common characteristics of the gut microbiome identified in patients with IBD are the reduction in bacterial diversity and development of a dysbiotic state, referring to alterations in the structure and function of the gut microbiome compared to healthy individuals [12, 14, 19]. Principal coordinate analysis with UniFrac [16] or Bray-Curtis [17] distance of the gut microbiome’s composition has identified differential clustering of healthy and IBD samples. Although the dysbiotic state is commonly identified in IBD patients, it remains unknown whether the microbiome initiates IBD or is only a reflection of the patient’s current health status. Larger meta-analyses have aimed to identify differentially abundant taxa between IBD patients and healthy controls in order to generate potential diagnostic biomarkers, although with limited success to date [18].

Due to difficulties identifying biomarkers with standard statistical methods for disease diagnosis, the field has moved to applying predictive machine learning (ML) models for classification of patient phenotypes. Several studies have demonstrated accurate classification of patients with IBD from their gut microbiome profile with ML models [2, 12, 13, 15, 18, 22–24]. Common ML models employed for IBD classification include random forest (collection of decision trees for classification) [2, 15], logistic regression (binary linear classifier) [13], and neural networks (layers of differently weighted nodes contributing to a classification) [23, 24].

Features commonly used for IBD classification with ML models can be categorized into three groups: clinical, bacterial, and functional. Clinical features encapsulate those regarding the patient (i.e. age, sex, body mass index (BMI)) and results from other clinical tests (i.e. calprotectin, colonoscopy), which are independent of a patient’s microbiome profile [25]. Taxonomy and functional features are usually determined via sequencing-based microbiome profiling, such as amplicon sequencing of the 16S rRNA gene or whole genome shotgun (WGS) sequencing of all DNA in a sample [26]. Bioinformatic tools, such as QIIME2 [27] or LotuS2 [28], provide pipelines for clustering 16S rRNA-amplicon sequences into operational taxonomic units (OTUs) which can then be compared to public databases to find taxonomy assignments [29]. WGS reads are frequently used to infer potential functions represented in the genomes of microbial community members (reviewed in [30]). Similarly, we can use known genomes in public databases to derive functional predictions in a community based solely on amplicon sequencing based taxonomy profiles, implemented in tools such as PICRUSt2 [31]. Although WGS provides greater taxonomic resolution and estimates of microbiome functions, 16S rRNA amplicon sequencing is currently more applicable to a diagnostic test due to its speed, affordability, and standardization of analysis tools.

A critical, and often under-explored, consideration for generating ML models for disease classification is their generalizability to previously unseen cohorts of patients. A ML model that underperforms when presented with data from a new patient cohort is not reliable enough to be applied in a clinical setting [32]. Despite this, models currently used in the context of microbiome data are often only trained and cross-validated with different splits of data from the same cohort. In studies where cross-validation with an unseen sample cohort is performed, the model’s performance is often lower, indicative of the model overfitting to the training set [22, 23]. A proposed explanation for the reduced performance is the potential for introduction of non-biological variability to the data by wet-lab protocols and sequencing instruments during the processing of these samples, typically observed in meta-analysis of microbiome data [12].

In order to improve model performance on unseen data, it is necessary to apply normalization and batch effect reduction techniques prior to model training. Normalization is a critical step to remove biases to feature abundance estimates, such as the data’s compositional nature, heteroskedasticity, or skewness. For example, microbiome data’s compositional nature prevents the direct application of standard statistical methods as they may lead to erroneous results, and requires prior application of compositional normalization methods [33, 34]. In addition, methods have been developed to remove the technical “batch effects” commonly identified in collections of samples from different studies, such as naive zero-centering methods and the recently developed empirical Bayes’ method, Meta-analysis Methods with a Uniform Pipeline for Heterogeneity (MMUPHin) [35–38]. To date, the effect of various combinations of normalization and batch effect reduction techniques on ML model generalizability remains to be benchmarked. In this article, we propose a standardized approach for evaluating the performance and generalizability of data processing pipelines and ML models with microbiome data to classify patients with IBD. Previous microbiome ML benchmarking studies focused on performance of various combinations of model type, normalization, and microbiome compositional features using variations of fivefold cross validation [24, 39]. Fivefold cross validation fails to assess the generalizability to new, unseen sample batches as each split potentially contains samples from all batches present in the dataset. Therefore, we implemented a leave-one-dataset-out (LODO) [40] cross-validation method to directly assess cross-batch generalizability. In this approach, the model is iteratively trained on samples of all but one dataset and then tested on the left-out dataset. Different combinations of data types, normalization methods, batch effect reduction methods, and ML models were assessed in order to establish a comprehensive performance benchmark of microbiome-based disease classification in the context of IBD.

## Results

### Overview of samples and methods

In order to assess the cross-batch performance of each pipeline, we implemented a LODO cross validation approach. We collected 16S rRNA gene next generation sequencing data from 15 studies in North America for a total of 7707 samples, comprising 55% healthy and 45% IBD samples, of which 56% are CD and 44% are UC **(Table 1)**. We completed 15 cross-validation iterations with a single dataset removed from the training set for generation of the classification model which was then used to assess model performance (**Figure 1)**.

**Figure 1.**
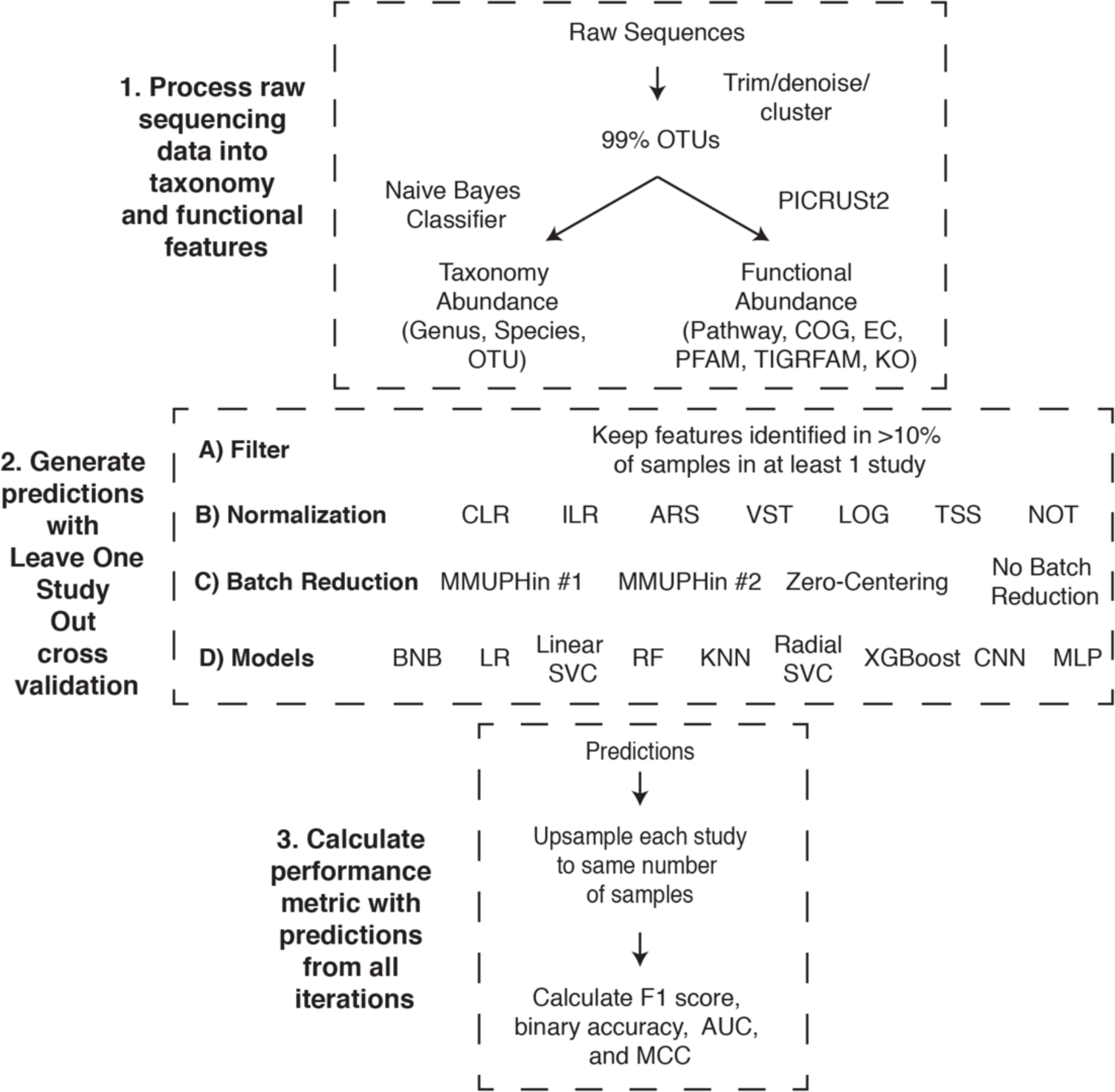
Leave-one-dataset-out cross-validation pipeline. The experiments comprised three different stages to go from raw sequence files to the performance metrics. 1) Raw sequences were processed with Dada2 or Deblur and close-reference clustered into OTUs at 99% identity. The OTUs were classified to taxonomy at 99% identity with QIIME2 and used to infer functional profiles with PICRUSt2. 2) Generating predictions for the 15 iterations of our LODO cross validation consisted of all possible combinations of the listed filtering method, normalization methods, batch effect reduction methods, and models. 3) The predictions from each iteration were combined and the number of samples from each dataset up sampled to 100 000 prior to calculating the performance metrics. The descriptions of acronyms and abbreviations are the following: Clusters of Orthologous Groups of proteins (COG), Kyoto Encyclopedia of Genes and Genomes (KEGG) orthologs (KO), Enzyme Commission (EC), Pfam protein domain (PFAM), TIGR protein family (TIGRFAM) and MetaCyc pathways (pathway), centered log-ratio (CLR), isometric log-ratio (ILR), arcsine square root transformation (ARS), variance stabilizing transformation (VST), log transformation (LOG), total sum scaling (TSS), no normalization (NOT), Bernoulli Naive Bayes (BNB), logistic regression (LR), linear support vector machine (Linear SVC), random forest (RF), K nearest neighbours (KNN), radial support vector machine (Radial SVC), eXtreme Gradient Boosting (XGBoost), convolutional neural network (CNN), multilayer perceptron (MLP).

**Table 1.**
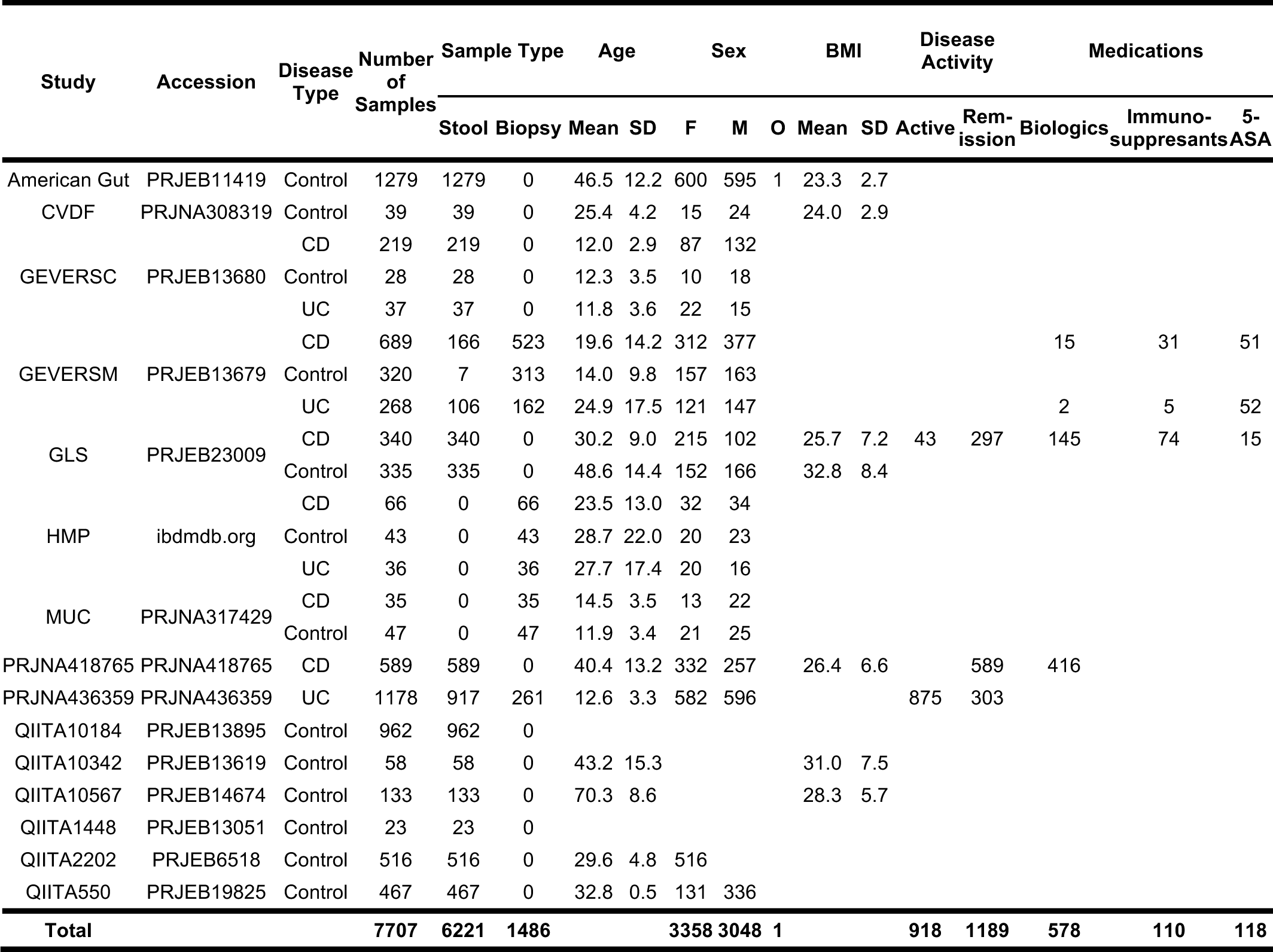
Overview of 15 datasets used to compare the effect of different features, data preprocessing methods, and machine learning models on IBD classification performance. Available metadata (age, sex, BMI), disease activity, and medication use) is provided for each dataset. Blank spaces indicate that the respective metadata was not available for the dataset’s samples. The following abbreviations are used: female (F), male (M), and other (O).

We evaluated the ability to classify samples from patients with IBD or non-IBD controls using different combinations of three taxonomic feature sets or six functional feature sets, eight normalization methods, four batch effect reduction methods, and nine machine learning models (**Figure 1**). The binary classification performance of each combination of feature set, normalization, batch effect reduction, and machine learning model was assessed with four classification metrics: F1 score, Matthews Correlation Coefficient (MCC), binary accuracy, and Area Under the receiver operating characteristics Curve (ROC-AUC, abbr. AUC) [60, 61]. We assessed generalizability through two methods. First, we sorted the pipeline components of interest (e.g. types of machine learning models) by the mean and standard deviation of their performance assessed by each metric. Second, in order to determine if the performance was significantly different, we performed statistical comparison of the pipelines’ metrics with a Mann-Whitney U test. Therefore, the most generalizable component was defined as the top sorted method which displayed significantly better performance than baseline or other methods.

### Top IBD classification was obtained using taxonomic features

Taxonomic features (species, genus or OTU) are predominantly used as input for ML models, whereas it is less common to use inferred functional features from PICRUSt2 as input. However, previous studies have identified lower inter-individual variation of the gut microbiome’s inferred functional profile than taxonomy [62, 63], suggesting that functional features may lead to better classification performance and generalizability. We processed the 16S sequencing samples with QIIME2 and PICRUSt2 to obtain taxonomy and functional feature abundance estimates, respectively.

For each ML model, we assessed the performance with taxonomy and functional abundance features in combination with normalization and batch effect reduction methods. Independent sorting of four classification performance metrics indicated that the taxonomic features classified IBD samples more effectively than functional features **(Figure 2A, Supplemental Figure 1A)**. Comparison of performance with taxonomy and functional features confirmed the significantly higher performance for classification of IBD samples with taxonomic features (**Figure 2B, Supplemental Figure 1B)**. Therefore, ML models using taxonomic features from this dataset lead to better classification of IBD samples than functional features.

**Figure 2.**
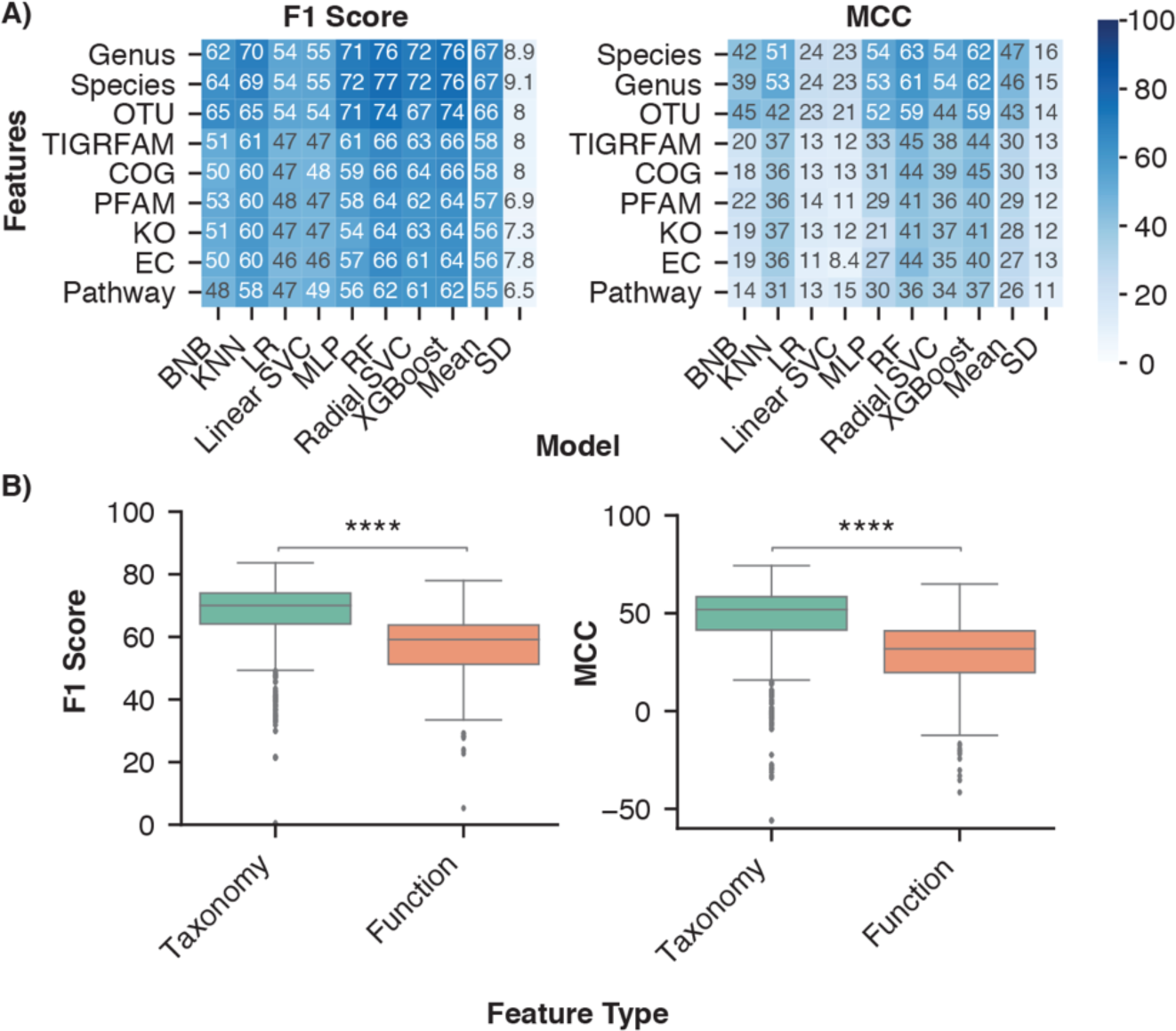
Optimal disease classification of microbiome samples obtained with taxonomic features. A) Average performance of the taxonomy and functional feature sets for each ML model architecture. Rows were sorted in descending order by the mean column followed by the standard deviation (SD) column. B) Distribution of performance metrics for taxonomy and functional features across all normalization, batch effect reduction, and model combinations. Independent Mann-Whitney U tests were performed to compare aggregate performance of taxonomy and functional features. The analysis was limited to normalization (ILR, CLR, VST, ARS, LOG, TSS, NOT) and batch effect reduction (no batch reduction or Zero-Centering) methods that were performed on all feature sets. **** indicates p-value < 0.0001.

Taxonomic classification with QIIME2 consists of seven hierarchical ranks, with kingdom and species at the top and bottom, respectively. Each consecutively lower taxonomy rank provides greater resolution of the gut microbiome’s composition while also increasing data sparsity, which can negatively affect an ML model’s performance [64]. Previous literature comparing different taxonomy ranks for disease classification indicated that lower ranks, down to genus, improved performance [65]. We assessed whether the trend for improved classification continued with the species rank and OTUs, despite their increasing sparsity. While no significant performance difference was observed between species and genus ranks, both displayed significantly higher classification performance than OTU features (**Supplemental Figure 1C)**.

### Non-linear models achieve greatest classification performance

Machine learning classification models identify decision boundaries within the feature space to separate sample labels from one another. For some ML models (BNB, Linear SVC, LR), these boundaries are linearly constrained, whereas others (RF, KNN, MLP, Radial SVC, XGBoost) can identify more complex, non-linear relationships between features and class. We assessed the generalizability of three linear and five non-linear ML models across the taxonomy and functional feature sets.

Independent sorting of ML models for each performance metric indicated that the non-linear models had greater classification performance (top five were non-linear models) than the linear models **(Figure 3A, Supplemental Figure 2A)**. Comparison of aggregate scores further confirmed non-linear models had significantly higher F1 score, MCC, AUC, and accuracy than linear models (**Figure 3B, Supplemental Figure 2B)**. In order to directly assess whether the non-linearity of a model improves classification in the context of microbiome data, we compared linear and non-linear variations of a support vector machine and logistic regression. Comparison of the two variations enables direct analysis of the impact of decision boundary constraints on performance, independent of differences in model architecture. The non-linear (radial) version of logistic regression and support vector machines (Radial) had significantly greater performance than the linear version (Linear) across all four metrics (**Figure 3C, Supplemental Figure 2C)**. Lastly, we assessed which non-linear model led to the highest classification performance. Across all four metrics, the random forest and XGBoost models were significantly better than MLP, KNN, and radial SVC models (**Supplemental Figure 2D)**. In conclusion, non-linear models provided more accurate IBD classification, likely due to the complex relationships between features and disease labels.

**Figure 3.**
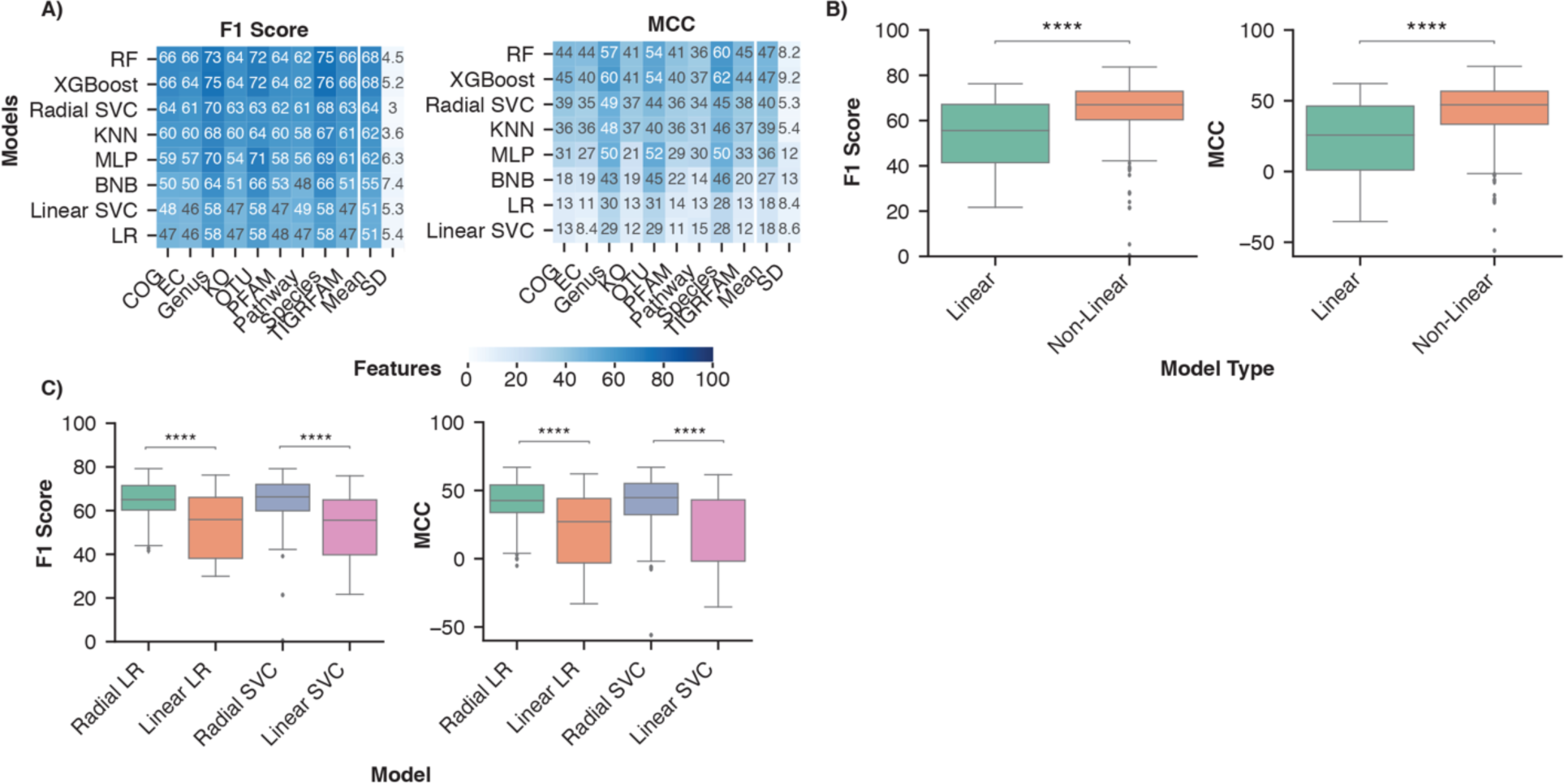
Non-linear models are better suited to identify decision boundaries between control and IBD samples than linear models. A) Average model performance for each feature set across normalization and batch effect reduction methods. Rows were sorted in descending order by mean followed by the standard deviation of performance across all feature sets. B) Distribution of performance of non-linear (RF, MLP, KNN, XGBoost, radial SVC) and linear (BNB, Linear SVC, LR) models. Independent Mann-Whitney U-tests were performed to compare each performance metric. The analysis was limited to normalization (ILR, CLR, VST, ARS, LOG, TSS, NOT) and batch effect reduction (no batch effect reduction or zero centering) methods performed on all feature types. C) Distribution of classification performance with the non-linear and linear variations of logistic regression and support vector machines across all feature sets. A Mann-Whitney U test with Bonferroni correction was performed to compare the linear and non-linear variation of each model respectively. **** indicates p-value < 0.0001.

Other ML model architectures, such as convolutional neural networks (CNNs), are commonly used for classification problems with certain structure in the input data, such as image classification. In the context of microbiome data, the CNN MDeep adds structure to OTU features through hierarchical agglomerative clustering of the phylogeny-induced correlation between OTUs [58]. As MDeep is currently only developed for OTU features, we assessed whether this CNN architecture led to greater classification performance with OTU abundance than our MLP architecture. Comparison of each performance metric across all normalization and batch effect reduction methods indicated MDeep performance was not significantly different than our MLP model (**Supplemental Figure 2E**).

Due to the significantly better performance of non-linear classification models and taxonomic features, our subsequent analysis of normalization and batch effect reduction methods utilized only taxonomic feature sets and non-linear models.

### Evaluation of normalization methods

We assessed normalization methods which account for different biases commonly observed in next-generation sequencing data: compositionality, heteroskedasticity, and skewness. We selected two normalizations designed for compositional data: the isometric log ratio (ILR) and centered log ratio (CLR) [66]. We selected two normalization methods which aim to reduce the heteroskedasticity: the arcsine square root (ARS) transformation [67] of the total sum scaling (TSS) values and the variance stabilized transformation (VST) from the R package DESeq2 [68]. Next, we assessed a log transformation of the TSS values (LOG), which reduces the positive skew commonly seen in the distribution of microbiome data. Lastly, we assessed normalization by TSS alone to remove differences in sequencing depth between samples or no normalization (NOT).

Independent sorting of each performance metric consistently identified the compositional normalization methods (CLR and ILR) as the most generalizable across non-linear models, followed by the variance/distribution modifiers (ARS, LOG, VST), and TSS as the consistently lowest performing normalization (**Figure 4, Supplemental Figure 3A**). Furthermore, the compositional methods led to significantly better performance than the other normalization types across all four metrics (**Supplemental Figure 3B**), whereas the variance/distribution modifiers and scaling method were only significantly better than no normalization. These results indicate the importance of normalization methods which account for the compositional properties of microbiome data prior to model training.

**Figure 4.**
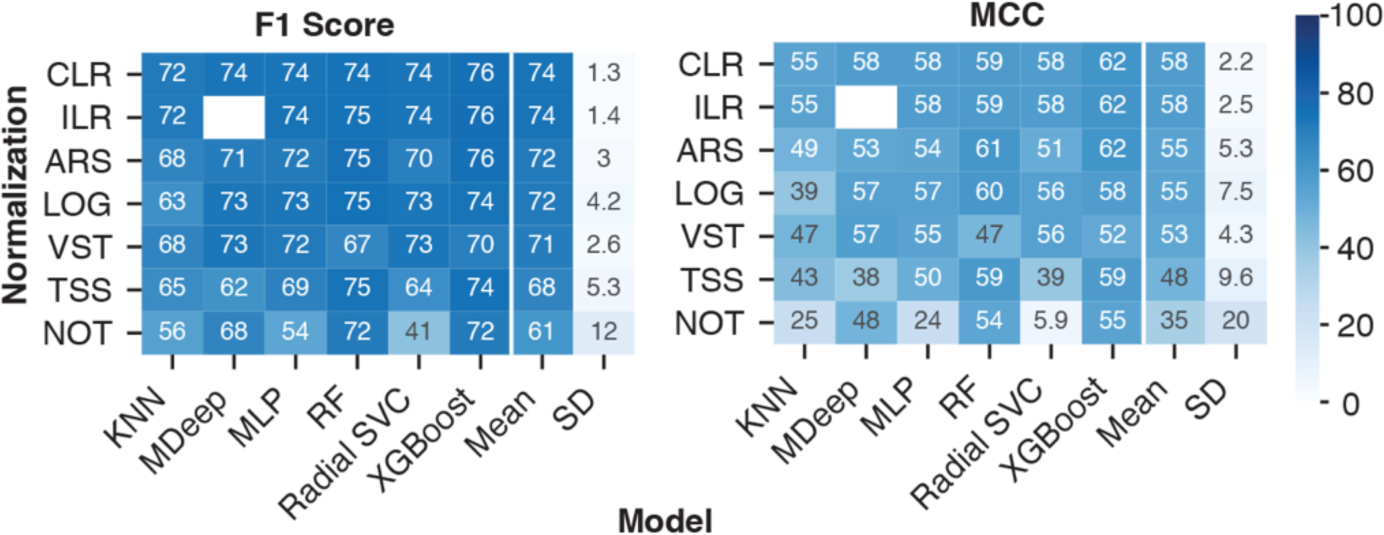
Compositional normalization methods lead to the highest model performance for IBD classification. Average model performance with each normalization method across all batch effect reduction methods. Performance following data processing with all pairwise combinations of the normalization methods (ILR, CLR, LOG, ARS, VST, TSS and NOT) and batch effect reduction methods (No batch reduction, MMUPHin #1, MMUPHin #2, and Zero-Centering) were included. Rows were sorted in descending order by the mean and standard deviation of each performance metric across the non-linear models. No analysis was performed for MDeep paired with ILR as the ILR normalized values no longer map directly to a feature, therefore removing the phylogenetic structure required for MDeep.

### Evaluation of batch effect reduction methods

A common issue with combining next-generation sequencing datasets for meta-analyses is the systematic differences between datasets due to differences in technical protocols. These differences add non-biological variation to the samples, decreasing the ability to ascertain biological signals [69]. Various approaches have been proposed to remove technical artifacts from dataset collections, of which we selected two relevant to microbiome data [37, 38]. First, zero-centering methods aim to reduce batch effects by centering the mean of each feature within a batch to zero. Second, Meta-analysis Methods with a Uniform Pipeline for Heterogeneity in microbiome studies (MMUPHin) [39] (microbiome specific empirical Bayes’ methods), estimate and remove batch-specific parameters for each feature. Two variations of MMUPHin were implemented to simulate the scenario of obtaining a new dataset when implemented for a diagnostic test. The first (#1) applied MMUPHin to the training and test sets separately, whereas the second (#2) only applied MMUPHin to the training set (see Methods for detailed description).

The different batch effect reduction methods were sorted by the mean and standard deviation of their performance across taxonomic features, non-linear models, and all normalization methods. Our sorting method indicated that zero-centering was the most generalizable approach across the non-linear models. Whereas MMUPHin #1 and #2 were less generalizable than no batch reduction, with MMUPHin #2 the least generalizable (**Figure 5, Supplemental Figure 4A**). Additionally, zero-centering led to significantly higher binary accuracy and MCC compared to all other methods, whereas F1 score and AUC were higher only when compared to no batch effect reduction and MMUPHin #2. MMUPHin #1 had significantly better performance than MMUPHin #2, with no difference in performance observed compared to no batch effect reduction (**Supplemental Figure 4B)**.

**Figure 5.**
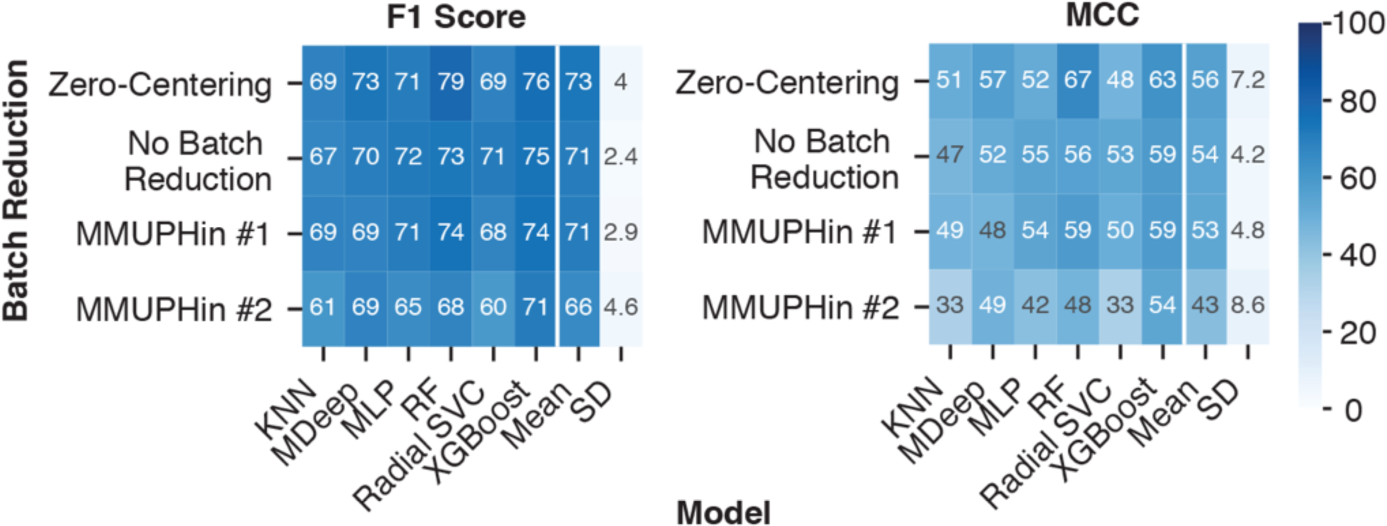
Batch effect reduction with the naive zero-centering method improved IBD classification. Average performance of each batch effect reduction method across all combinations of normalization methods, taxonomic features, and non-linear ML models. Performance following data processing with all pairwise combinations of the normalization methods (ILR, CLR, LOG, ARS, VST, TSS and NOT) and batch effect reduction methods (No batch reduction, MMUPHin #1, MMUPHin #2, and Zero-Centering) were included. Rows were sorted in descending order by the mean and standard deviation of performance across all non-linear models.

### Evaluation of model performance on sample and patient subgroups

The samples used to assess the performance of different combinations of normalizations, batch effect reduction, and ML models were drawn from across sample collection methods (i.e. stool and biopsy) and patient demographics (i.e. paediatric and adult samples). While we did not set inclusion criteria for samples based on these differences, previous research has demonstrated distinct differences in microbiome composition between sample types and demographic groups [18, 70, 71]. For example, principal coordinate analysis (PCoA) with weighted UniFrac distance [72] and principal component analysis (PCA) of CLR-transformed taxonomic features indicated paired biopsy and stool samples from the same individual cluster separately [73].

We compared the model performance for the sample and patient demographics for which we were able to acquire sufficient metadata and have been associated with microbiome alterations: sample type (biopsy vs. stool), IBD subtype (CD vs. UC), sex (Female vs. Male), BMI (BMI < 30 vs. BMI > 30), and age (Adult vs. Pediatric). To assess the performance within each demographic, we included the predictions from taxonomic features (species, genus, OTU) with a compositional normalization method, zero-centering batch effect reduction, and a non-linear ML model. Our analysis focused on the MCC performance metric as it is more robust to imbalanced label distribution [61], which occurred when the samples were grouped by the five metadata categories mentioned. A logistic regression function was used to assess changes in performance corresponding to each demographic while controlling for the other metadata (**Table 2)**.

**Table 2.** Model performance for different sample types and patient demographics. Samples with available metadata were categorized into groups based on the collection method or the patient’s specific demographic group based on sex, age, and BMI. Predictive performance for all combinations of taxonomic features, compositional normalizations, zero-centering batch effect reduction, and non-linear models were included in the analysis. Logistic regression was performed to assess the performance differences within each sample and demographic group while adjusting for the remaining covariates. **** indicates p-value < 0.0001, and * indicates p-value < 0.05. Coefficient refers to the corresponding independent variable’s coefficient for the logistic regression function and SE refers to the standard error of the coefficient.

| Group | Variable | Coefficient | SE |
| --- | --- | --- | --- |
| Sample Type | Biopsy (vs. Stool) | -0.44 * | 0.2 |
| Life Stage | Adult (vs. Pediatric) | 1.39 **** | 0.18 |
| BMI Stratification | BMI <30 (vs. BMI > 30) | -0.85 **** | 0.19 |
| Sex | Female (vs. Male) | -0.02 | 0.18 |
| IBD Type | CD (vs. UC) | 0.05 | 0.18 |

The models displayed reduced performance for biopsy samples compared to stool samples, increased performance for samples from adult patients compared to paediatric patients, and decreased performance of samples from patients with BMI less than 30 compared to patients with BMI greater than 30. On the other hand, there was no difference in classification performance for females compared to males or for samples from patients with CD compared to patients with UC (**Table 2**). Similar results were reproduced with F1 scores, AUC, and accuracy (**Supplemental Table 1**). The metadata groups with different performance between the two categories coincided with those that are not equally represented in our dataset, highlighting the importance of accounting for different demographic groups in a microbiome based diagnostic test.

### Evaluation of top performing pipeline combinations for IBD classification

Our analysis identified the features, ML models, normalization methods, and batch effect reduction methods which led to the most generalizable performance. In order to determine the best overall combination of features, data processing, and ML model we assessed the top three performing models (**Table 3**). The top three models consisted of the most generalizable individual components: taxonomic features (genus), non-linear model (XGBoost or RF), compositional normalization (ILR or CLR), and zero-centering to remove batch effects. Therefore, the combination of the most generalizable methods led to the best classification performance.

**Table 3.** Top three data processing and model pipelines for classifying IBD samples. Three combinations which appeared most frequently when all models were sorted by F1 score, accuracy, AUC, or MCC.

| Features | Normalization | Batch Reduction | Model | F1 Score | Accuracy | AUC | MCC |
| --- | --- | --- | --- | --- | --- | --- | --- |
| Genus | ILR | Zero-Centering | XGBoost | 83.67 | 87.92 | 87.9 | 74.3 |
| Genus | CLR | Zero-Centering | XGBoost | 82.96 | 87.41 | 87.3 | 73.2 |
| Genus | ILR | Zero-Centering | Random Forest | 82.66 | 87.4 | 86.9 | 72.9 |

### Identification of important features for classification with a XGBoost model

In addition to predicting disease diagnoses, machine learning models can be used to identify biomarkers for disease by identifying features important for disease classification. We characterized the feature importance from the second-best overall data processing and ML model pipeline (**Table 3**). We did not analyze the feature importance of the best-performing model because the ILR normalized values no longer correspond to the starting features thereby preventing interpretation of feature importance. For an XGBoost model, the importance corresponds to a feature’s contribution to the model’s decision during training, referred to as the gain value [57]. We extracted the features’ gain values from each of the 15 LODO iterations, sorted by the mean of all iterations, and plotted the top fifteen features (**Figure 6)**. In addition, we determined the change in abundance for each taxonomy to assess whether our dataset aligned with previous findings on changes of the microbiome in IBD.

**Figure 6.**
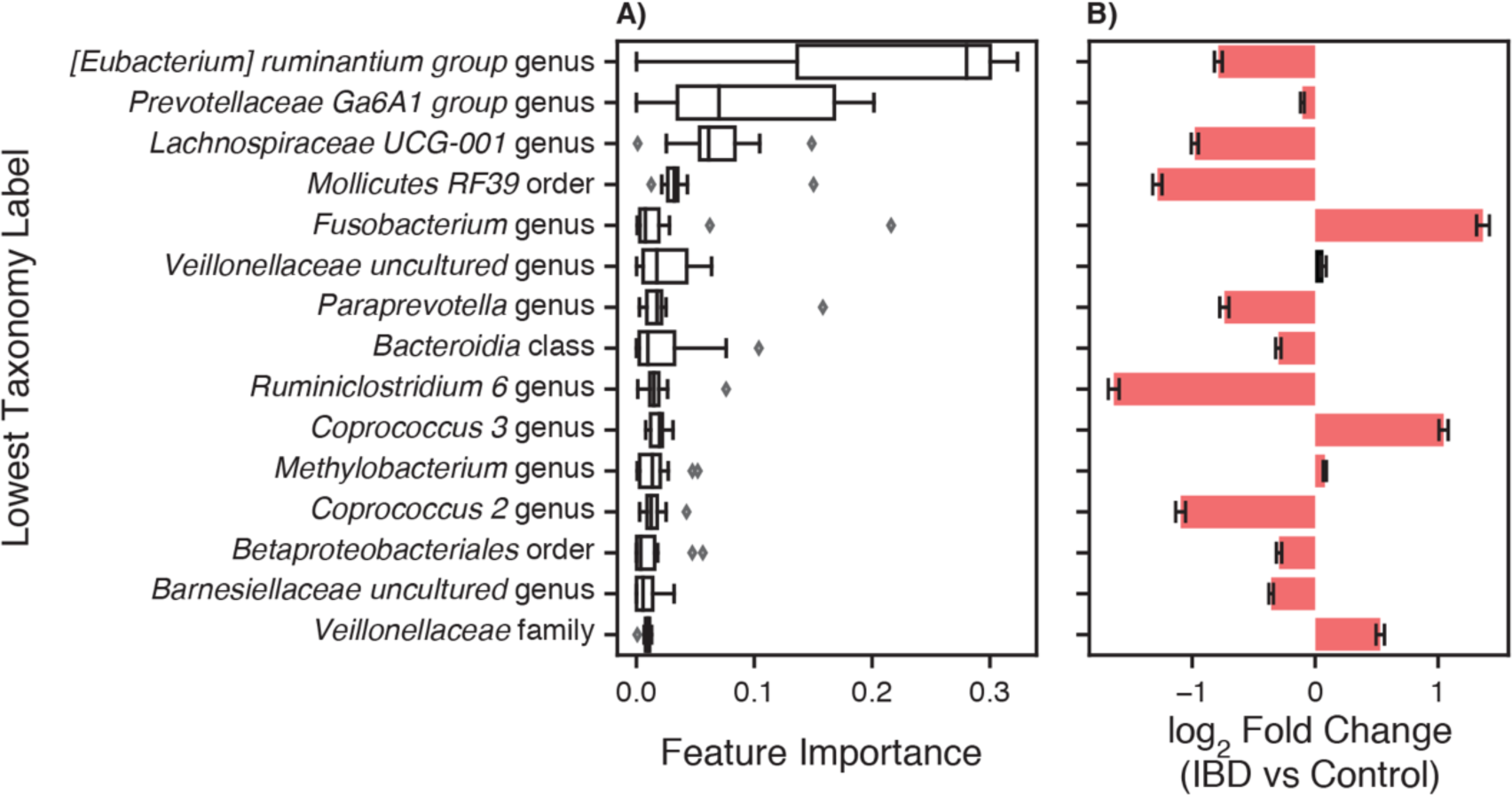
Features with greatest contribution to IBD classification by an XGBoost classifier. A) An XGBoost classifier was trained with CLR normalized genus abundance features and zero-centered batch effect reduction for fifteen LODO iterations. The features’ gain values for each iteration were extracted and sorted by the mean gain across all iterations. The lowest classification rank for each feature was used as the label for the corresponding bar. B) Changes in taxonomy abundance between control samples and those from patients with IBD. Bars represent the fold change ± the standard error determined with Analysis of Compositions of Microbiomes with Bias Correction (ANCOM-BC). Red indicates a significant fold change between IBD and control samples (p < 0.05) and black indicates non-significant fold change.

Amongst the top features are many taxa in the short chain fatty acid (SCFA) producing Clostridium XIVa/IV clusters, including bacteria from the *Eubacterium*, *Coprococcus*, *Lachnospira*, and *Ruminiclostridium* genera (**Figure 6A)**. Aligning with previous studies, these bacteria were decreased, with the exception of *Coprococcus 3*, in IBD samples vs control samples in our dataset (**Figure 6B)** [2, 74]. *Fusobacterium* and *Veillonellaceae* genera, commonly increased in the gut microbiome of IBD patients, were also top contributors to the XGBoost classifier (**Figure 6A/B)** [2, 75]. In addition, the *Prevotellaceae* genus was the second most important feature, with the decreased abundance in IBD samples agreeing with previous studies showing decreased abundance in the gut microbiome of patients with CD and UC (**Figure 6A/B)** [76]. XGBoost classifiers have the best potential for use as a diagnostic test due to their performance as well as their interpretability and utility in identifying disease biomarkers.

## Discussion

We assessed how different feature sets, ML models, normalization methods, and batch effect reduction methods affect predictive performance across patient cohorts in a LODO cross validation approach. The limited applicability of a PCR-based diagnostic test with a handful of microbiomes for IBD diagnosis [77] has led the field to explore the use of ML models for disease diagnosis. Our benchmark provides practical suggestions for ways to improve the performance of an IBD diagnostic test using the gut microbiome composition. First, genus abundance estimates from 16S rRNA sequencing need to be normalized by a compositional normalization method, with CLR normalization being the most appropriate as it allows for each features importance to the ML models decision to be assessed. Second, zero-centering batch effect reduction should be applied to each batch of samples collected, sequenced, and processed together to reduce systematic batch differences. Following normalization and batch effect reduction, an XGBoost or random forest classification model should be trained and optimal hyperparameters determined for implementation as a diagnostic test. With respect to the training dataset, it is important to account for patient demographics or technical differences between samples that have been associated with gut microbiome alterations. We suggest several options for optimal performance: (1) ensure balanced representation in the training dataset, (2) include the metadata labels as a feature for the model, or (3) deploy diagnostic ML models built specifically for one demographic group. In addition, the LODO cross-validation methodology is an important tool for the selection of new data preprocessing and model building methods.

Previous studies have demonstrated greater consistency of functional feature abundances than taxonomic feature abundance in both healthy individuals [78–80] and those with IBD [63, 81]. In fact, some studies were unable to identify a single bacterium present in every IBD patient from their cohort [62]. The reduced variation and sparsity of functional features led us to hypothesize that functional abundance profiles would lead to better classification of IBD samples. However, through our LODO cross validation, we found that classification performance with functional features was significantly worse than with taxonomic features **(Figure 2B, Supplemental Figure 1B)**. We postulate the reason for the reduced classification performance with functional profiles is due to the limited recapitulation of functional profiles with PICRUSt2 [31, 82] and the inability of 16S rRNA sequencing to identify strain-level functional differences of the present bacteria [83]. To overcome these limitations in future studies, measurement of the microbiome’s gene content by WGS, transcriptomes by RNA-seq, or metabolites by metabolomics need to be explored. In fact, functional profiles from whole genome sequencing led to better predictions of patients with IBD who achieved remission with vedolizumab than taxonomy abundance [23]. While whole genome sequencing may improve disease classification, its much higher cost than 16S rRNA sequencing substantially hinders the technology’s adoption as a diagnostic test.

A major hurdle in the implementation of sequencing based diagnostic tests in the clinic is the observed systematic differences between sample preparations. In a previous study, removal of these batch effects with an empirical Bayes’ or zero centering approach led to improved classification [84]. However, our work only identified improved cross-batch classification performance with zero-centering and not the empirical Bayes’ method MMUPHin (**Figure 5)**. Current empirical Bayes’ approaches are designed and optimized for disease mechanism and biomarker discovery where the disease covariate is known and incorporated in the method. The inclusion of a disease covariate is not applicable to a diagnostic scenario though, where the diagnosis label is to be determined. The lack of improvement in classification performance with MMUPHin #1 compared to no batch reduction is potentially due to its implementation in a scenario the method was not optimized for.

Similar to batches of samples collected for a diagnostic test, the batches in our dataset were not balanced, with some containing only a single diagnosis class (e.g. all samples coming from IBD patients). In cases where the batch and diagnosis label are confounded, batch correction methods tend to reduce the disease associated differences in the process of removing the batch differences [37]. Therefore, the more advanced removal of batch effects by MMUPHin likely led to an over-adjustment within the unbalanced batches and removal of the disease differences. Whereas, the less sophisticated removal of batch effects with the covariate naive zero-centering approach retained sufficient biological signal between disease labels for non-linear ML models to correctly classify samples across batches. Batch correction methods that do not require input of a covariate have been developed, such as frozen surrogate variable analysis or reference principal component integration (RPCI) [85, 86], although their applicability to microbiome data has not been assessed.

The sparse availability of metadata for the samples led to several limitations in our analysis. First, the identification of CD and UC patients relied on the accuracy of the diagnosis coding in the public databases. However, there were no studies explicitly validating the registration of CD and UC diagnosis codes. Second, although our study demonstrated reliable results, gaps in the publicly available data prevented us from several critical analyses. For instance, we lacked information on how the patients were diagnosed in every study, the timing of sample collection in relationship to their diagnosis and disease progression, current disease activity quantification, DNA extraction and sample storage information. Furthermore, there was limited information on environmental factors such as medication usage, alcohol usage, smoking, diet, and other factors known to alter the gut microbiome which could affect our analysis [81, 87]. Of the sample information and patient demographic data we obtained, clear differences in performance of our top pipelines were observed (**Table 2**). Therefore, future studies with improved lifestyle and clinical metadata are needed to systematically address how these factors affect performance of a gut microbiome diagnostic test.

Other non-invasive diagnostic tests for IBD, such as fecal calprotectin, continue to have significant differences between the reports on the sensitivity and specificity for classifying IBD patients from non-IBD [88, 89]. While high performance levels have been reported, one recent study identified a 78% accuracy for identifying patients with IBD using fecal calprotectin [90], which is approximately 10% lower than our best model. Furthermore, while we focused solely on IBD classification, ML models using microbiome composition have wider applicability than singular biomarkers such as calprotectin. Models using microbiome data have already been implemented to predict if a patient with IBD will respond to a medication [23], to predict a patient’s postprandial glycemic response [91], and for classification of other diseases, such as Parkinson’s disease [48], to name a few.

## Conclusion

With sufficient data and validation, analysis of the fecal gut microbiome can indeed be leveraged as a multi-purpose predictive tool. Given the significant delay [92–94] and associated costs of diagnosis [95, 96], it is critical to continue exploration of approaches that increase accessibility of diagnosis and decrease the cost of testing [97] in a community health or primary care setting. XGBoost and random forest machine learning models with microbiome data have the potential to achieve these goals. Further work to gather more well-annotated data, improve performance and assess models with validation studies is required.

## Methods

### Acquisition of sample data

Sample FASTQ files were acquired from the European Nucleotide Archive (ENA) browser. The sample metadata was acquired from the corresponding publication’s supplementary materials or the QIITA microbiome platform. Only samples collected from individuals in North America were used from each dataset. The dataset accessions and technical information regarding the samples in each dataset are available in **Supplemental Table 2**.

The following fifteen studies were included in our dataset:

1. The American Gut cohort is from a large, open platform which collected samples from individuals in the US to identify associations between microbiomes, the environment, and individual’s phenotype [41]. We included available samples that did not contain any self-reported diseases in the metadata.
2. The CVDF study determined the effect of cardiorespiratory fitness on microbiome composition and comprises a range of fitness levels [42].
3. The GEVERSM study assessed the microbiome composition of treatment naive, newly diagnosed, paediatric patients with IBD and adult patients diagnosed with IBD for 0 to 57 years [2].
4. The GEVERSC cohort consists of additional samples from paediatric and adult patients added to the GEVERSM study [2].
5. The GLS study longitudinally sampled 19 patients with CD (Crohn’s disease activity index (CDAI) between 44 and 273) and 12 healthy control individuals [20].
6. The Human Microbiome Project (HMP) study longitudinal tracked paediatric and adult patients ranging from newly diagnosed to diagnosed for 39 years. Diagnosis was confirmed by colonoscopy prior to enrollment in the study along with several other inclusion criteria listed in the corresponding publication [43].
7. The MUC study collected mucosal biopsies from 44 paediatric patients with CD and 62 non-IBD paediatric control patients [21].
8. PRJNA418765 was a longitudinal study of patients with CD that were refractory to anti-TNF initiating ustekinumab assessed at week 0, 4, 6 and 22. To be included, patients required at least three months Crohn’s disease history and a CDAI between 220 and 450 [44].
9. PRJNA436359 was a longitudinal study of new onset and treatment naive paediatric patients with UC receiving a variety of medications at week 0, 4, 12, and 52. Inclusion criteria consisted of presence of disease beyond the rectum, Paediatric Ulcerative Colitis Activity Index (PUCAI) of 10 or more, and no previous therapy [45].
10. QIITA10184 was a study comparing five different fecal collection methods and their effect on the healthy participant’s microbiome composition identified with 16S rRNA gene sequencing [46].
11. QIITA10342 study assessed the microbiome composition and function of healthy individuals in two American Indian communities in the United States [47].
12. QIITA10567 samples consist of the control individuals in a study linking alterations in microbiome composition to Parkinson’s disease [48].
13. The QIITA1448 study compared microbiome composition of individuals in traditional agricultural societies in Peru to those in industrialized cities in the United States [49].
14. The QIITA2202 study collected longitudinal stool samples from two healthy individuals alongside detailed lifestyle characteristics to correlate with microbiome composition [48, 50].
15. The QIITA550 study collected longitudinal stool samples from two individuals to assess temporal changes in microbiome composition [51].

### Taxonomy classification with QIIME2

Taxonomy abundance tables were generated from the FASTQ files using QIIME2 (v2020.2) [27]. Reads were trimmed to remove low quality reads (trimming parameters listed in Supplemental Table 1), chimeras removed, and sequences denoised using Dada2 [52] or Deblur (for GLS and AG only). The processed sequences were clustered into OTUs and the centroid sequences classified with a Naive Bayes classifier [53] at 99% identity using the Silva 132 99% reference database [29, 54, 55]. For classification, the corresponding 16S rRNA gene hypervariable region’s sequences were extracted from the Silva 132 99% reference database with the QIIME2 plugin feature-classifier’s extract-reads function using the primers from the respective study. The extracted reads and the corresponding taxonomy were used to train the Naive Bayes classifier with the QIIME2 plugin feature-classifier’s fit-classifier-naive-bayes function. Taxonomic feature tables were collapsed to species (level 7) and genus (level 6) classification for further analysis.

### Inferring Functional Abundance with PICRUSt2

Functional abundance tables were generated using PICRUSt2 (v2.3.0) from the OTU abundance table and representative OTU sequences from QIIME2. We generated abundance tables from the six different databases incorporated into PICRUSt2: Clusters of Orthologous Groups of proteins (COG), Kyoto Encyclopedia of Genes and Genomes (KEGG) orthologs (KO), Enzyme Commission (EC), Pfam protein domain (PFAM), TIGR protein family (TIGRFAM) and MetaCyc pathways. Each database is independently curated and provides information on different aspects of the functional properties present in the microbiome.

### Leave-One-Dataset-Out (LODO) cross validation

The generalizability of each model, normalization, and batch effect reduction method, was determined through a cross validation strategy which assessed predictive performance on previously unseen batches of samples **(Supplemental Figure 5)**. As there were 15 datasets, we iterated through the full dataset 15 times, generating the training set by removing all samples from a single dataset to a separate test set. The training set was used to prune features that were not present in at least 10% of samples from one dataset. Following pruning, the remaining features were selected from the test set and the samples were normalized and batch reduced with the respective methods. Lastly, the training set was balanced to have the same number of healthy and IBD samples by subsampling the label with the greater number of samples, while maintaining the proportion of samples from each collection site, disease label (UC/CD/Control), and sample type (stool/biopsy).

For our modified implementation of MMUPHin, the data processing was adjusted to ensure the training and test sets were batch reduced separately. For MMUPHin #1, the test dataset’s samples were removed and the data from the remaining studies batch reduced with MMUPHin prior to training the model. Independently, the full dataset (with training studies and test dataset) was batch reduced and the test dataset’s samples then used to assess the model’s classification performance (**Supplemental Figure 5**, MMUPHin #1). For MMUPHin #2, the training studies were batched reduced with MMUPHin prior to model training and the model’s classification performance then assessed on non-batch reduced samples from the test dataset (**Supplemental Figure 5**, MMUPHin #2). Lastly, feature abundance for some samples following MMUPHin batch effect reduction on the training set when QIITA2202 was left out and the test set when HMP was left out were all zero. Rows with all zero are not appropriate input for the compositional normalization methods, therefore we replaced the feature values for these samples with equal relative abundance prior to normalization.

### Feature Selection

Following taxonomy classification and inference of functional abundance, features present in less than 10% of the samples within each dataset were pruned from the dataset. The feature pruning was performed on the training set only, with the features then selected from the test set.

### Normalization methods

When possible, normalization methods were implemented using python (v3.6.12) and R (v3.6.3) packages with the methods already incorporated. For CLR and ILR normalization, zero values were first replaced with the multiplicative replacement function prior to normalization with the clr and ilr functions, respectively, from the python package SciKit-Bio (v0.5.2). CLR performs a log transformation of abundance values, which are normalized by the geometric mean of all features. ILR uses a change of coordinate space projection calculation to transform proportional data (or relative abundances) to a new space with an orthonormal basis.

For TSS normalization, the counts for each feature were divided by the sum of all feature counts in the sample with a custom python function. The method constrains the sample row sum to one, aiming to similarly scale all samples while maintaining biological information of microbial abundances. For ARS normalization, the TSS normalized values were transformed with sqrt function followed by the arcsin function from the python package numpy (v1.19.2). The LOG normalization was also applied to the TSS normalized values using the log function from numpy following replacement of all 0s with 1.

For VST normalization, we used the varianceStabilizingTransformation function in the R package DESeq2 (v1.26.0). VST aims to factor out the dependence of the variance in the mean abundance of a feature. The method numerically integrates the dispersion relation of the feature mean fitted with a spline, evaluating the transformation for each abundance in the feature. VST normalization was performed similarly to the previously described modified MMUPHin implementation, with the training set normalized separately from the test set as the normalization is dependent on all samples present in the dataset.

### Batch effect reduction methods

We explored two methods for batch effect reduction: naive zero-centering and an empirical Bayes method. The naive zero-centering batch effect reduction entails centering the mean of each feature within each batch to zero [37]. We also assessed MMUPHin, a recently developed empirical Bayes method designed specifically for zero-inflated microbial abundance data. MMUPHin estimates parameters for the additive and multiplicative batch effects, using normal and inverse gamma distributions, respectively. The estimated parameters are then used to remove the batch effects from the dataset [38, 56]. For MMUPHin, the sample type (stool/biopsy) was used as a covariate for MMUPHin #1 and the sample type and disease label (UC/CD/Control) were covariates for MMUPHin #2. We considered a batch as the whole dataset or split a dataset into multiple batches when the metadata indicated different sample preprocessing methods or samples were processed in different locations.

### Standard machine learning models

We assessed the classification performance of standard machine learning and deep learning models. The standard models were implemented using the python package SciKit-Learn (v0.23.2). Hyperparameters were not optimized and decided prior to experimentation.

#### Bernoulli Naive Bayes Classifier

The Bernoulli Naive Bayes Classifier (BNB) model converts the feature space to binary values and then estimates parameters of a Bernoulli distribution for classification purposes. We implemented the BNB model using the default settings in SciKit-Learn.

#### Random Forest

Random Forest (RF) models use an ensemble of decision trees that discriminate the feature space by a sequence of threshold conditional statements. The power of the model comes from its non-linear classification capabilities and the number of trees used to label classification. We implemented the Random Forest classifier with the following modifications to the default SciKit-learn settings: n_estimaters = 500, max_features = sqrt, and class_weight = balanced.

#### K-Nearest Neighbour Classifier

The K-Nearest Neighbour Classifier (KNN) classifies each sample by majority vote of the K nearest neighbours in its surrounding. We implemented the K Neighbors classifier with the following modifications to the default SciKit-learn settings: n_neighbors = 6, weights = distance, and metric = manhattan.

#### Support Vector Machine Classifier

The Support Vector Machine Classifier (SVC) identifies multivariate decision boundaries that separate class labels. We implemented two SVC variations, the first with a linear kernel, constraining the decision boundary to a linear hyperplane, using the SGDClassifier class from SciKit-learn with the following modifications to default settings: loss = modified_huber, tol = 10e-5, and max_iter = 10000. The second variation used the radial basis function kernel with the SVC class from SciKit-Learn, which removes the linear constraint of the decision boundary, with the following modifications to the default settings: tol = 10e-6, class_weight = balanced, and max_iter = 100000.

#### Logistic Regression

Logistic Regression classification estimates the probability of a certain class in a binary classification problem using a statistical fit to the logistic function. We implemented the LogisticRegression class from SciKit-Learn with the following modifications to the default settings: solver = sag, class_weight = balanced, and max_iter = 10000. For the non-linear variation, the feature space was first transformed with the radial basis function kernel implemented with the rbf_kernel function from SciKit-Learn prior to fitting a logistic regression model.

#### Gradient Boosted Trees (XGBoost)

Gradient boosted trees consist of a collection of sequential decision trees, where each tree learns and reduces the error of the previous tree [57]. The gradient boosted trees model was implemented with the XGBoost package’s (v1.2.0) XGBoostClassifier class with the following modifications to default settings: n_estimators = 500.

### Deep learning models

The deep learning models were built with the python package Tensorflow (v2.2.0). The models were trained for up to 100 epochs with a batch size of 16 and samples shuffled. The best weights were selected using early stopping (EarlyStopping callback) by monitoring the validation loss (5% split of the training set) with a min_delta = 1×10^−3^ and patience = 10.

#### Multilayer Perceptron (MLP)

A MLP is a neural network architecture composed of one or more layers of fully connected neurons that take as input the weights of the previous layer and output the result of an activation function to the subsequent layer. For binary classification, the final layer contains a single node that predicts the class probability. We implemented an MLP architecture with three hidden layers of 256 neurons using a rectified linear unit (ReLU) activation function followed by a Dropout layer with a dropout rate of 50%. The final layer predicted the class label with a sigmoid activation function. The model was trained using a binary cross entropy loss function and the Adam optimizer with a learning rate of 0.001.

#### Convolutional Neural Network

We implemented MDeep, a CNN architecture recently designed for microbiome data [58]. CNNs require an inherent structure to present in the data, which is added to the OTU dataset by hierarchical agglomerative clustering of the phylogeny-induced correlation between OTUs. We built a phylogenetic tree with the align_to_tree_mafft_fasttree function in the QIIME2 phylogeny python plugin using the OTU representative sequences obtained from clustering 16S rRNA sequences with QIIME2. The phylogenetic tree was imported into R using the phyloseq package and the cophenetic distance between OTUs determined with the R package ape. The cophenetic distance was then used to calculate the phylogeny-induced correlation as described in the original study and OTUs clustered using the HAC function from the MDeep GitHub repository (https://github.com/lichen-lab/MDeep).

### Performance metrics

To measure the performance of the various normalization, batch effect reduction, and model combinations we used four commonly used metrics for binary classification: F1 score, Area Under the receiver operating characteristic Curve (AUC), binary accuracy and Matthews Correlation Coefficient (MCC). Since the number of samples in each dataset ranged from 23 to 1279, we first balanced the number of samples from each dataset by up sampling each (with replacement) to 100 000 samples while maintaining the confusion matrix proportions for each individual dataset. Balancing the number of samples ensured that altered performance with a single, large dataset did not control the overall score and changes in performance for small studies was still observed. The up sampled dataset was then used to calculate the respective metrics using the functions implemented in SciKit-Learn.

### Sample subgroup performance analysis

We assessed the performance of our algorithm for five different metadata variables, each with two categorical labels. The samples were grouped by the five variables, with the two categories for each variable coded as 0 or 1. The performance metric was calculated within each grouping for classification of control samples and either UC or CD (depending on the specific grouping). For the logistic regression analysis, the metric was input as the dependent variable and the five metadata groups as the independent variables. The MCC score was scaled with the MinMaxScaler from SciKit-Learn to scale the range from 0 to 1 as required for the logistic function.

### Feature Importance from XGBoost Classifier

In order to determine the importance of each taxonomy, we collected the features’ gain value from our second-best pipeline composed of CLR normalized, zero-centered, genus abundance features with an XGBoost Classifier. The gain values were collected from the trained XGBoost classifier in each LODO iteration separately.

### Taxonomy Differential Abundance

Differential taxonomy abundance was performed with Analysis of Compositions of Microbiomes with Bias Correction (ANCOM-BC) (v1.0.5) [59]. The fold change between control samples and IBD samples (UC and CD) was determined with a Bonferroni multiple comparison correction applied to the p-values.

## Abbreviations

ANCOM-BC: Analysis of Compositions of Microbiomes with Bias Correction
ARS: Arcsine square root transformation
AUC: Area Under the receiver operating characteristics Curve
BMI: Body mass index
BNB: Bernoulli Naive Bayes Classifier
CD: Crohn’s disease
CDAI: Crohn’s disease activity index
CLR: Centered-log ratio
CNN: Convolutional neural network
COG: Clusters of Orthologous Groups of proteins
EC: Enzyme Commission
HMP: Human Microbiome Project
IBD: Inflammatory Bowel Disease
ILR: Isometric-log ratio
KEGG: Kyoto Encyclopedia of Genes and Genomes
KNN: K-Nearest Neighbour Classifier
KO: KEGG orthologs
LODO: Leave-one-dataset-out
LOG: Log transformation
LR: Logistic Regression
MCC: Matthews Correlation Coefficient
ML: Machine learning
MLP: Multilayer Perceptron
MMUPHin: Meta-analysis Methods with a Uniform Pipeline for Heterogeneity
NOT: No normalization
OTU: Operational taxonomic unit
PCA: Principal component analysis
PCoA: Principal coordinate analysis
PFAM: Pfam protein domain
PUCAI: Paediatric Ulcerative Colitis Activity Index
RF: Random Forest
RPCI: Reference principal component integration
SCFA: Short chain fatty acid
SVC: Support Vector Machine Classifier
TSS: Total sum scaling
TIGRFAM: TIGR protein family
UC: Ulcerative Colitis
VST: Variance stabilized transformation
XGBoost: eXtreme Gradient Boosting
WGS: Whole genome shotgun

## Declarations

### Ethics approval and consent to participate

Not applicable.

### Consent for publication

Not applicable.

### Availability of data and material

Publicly available datasets were analyzed in this study. The raw sequencing data for the following 16S rRNA datasets were downloaded from European Nucleotide Archive at the following accession numbers: American Gut (PRJEB11419), CVDF (PRJNA308319), GEVERSC (PRJEB13680), GEVERSM (PRJEB13679), GLS (PRJEB23009), MUC (PRJNA317429), PRJNA418765, PRJNA436359, QIITA10184 (PRJEB13895), QIITA10342 (PRJEB13619), QIITA10567 (PRJEB14674), QIITA1448 (PRJEB13051), QIITA2202 (PRJEB6518), QIITA550 (PRJEB19825). The raw sequencing data for the HMP 16S rRNA dataset was downloaded from ibdmdb.org.

### Authors’ contributions

RK, JD, TZ, and RM designed the data processing pipeline, performed the experiments and analyzed the pipelines’ performance. RK and RM wrote the manuscript. AHG and SB contributed to the experimental design. AHG, SB, FH, TK, SK, PJ, and KK contributed to interpretation of the results and editing and revising the manuscript. All authors reviewed, revised, and approved the final manuscript.

## Acknowledgements

We would like to thank Luca Cuccia, Laura Minkova, Houman Farzin, Michael Golfi, Paul Godin, and Yasmine Mouley (Phyla Technologies Inc.) for their feedback and support as the manuscript was completed. We would also like to thank Sébastien Giguère (Valence Discovery) for his guidance during our methodology development.

## Author Information

**Phyla Technologies Inc, Montréal, Canada**

Ryszard Kubinski, Jean-Yves Kepaou Djamen, Timur Zhanabaev, Sani Karam, Kamran Kafi, Ryan D. Martin

**Mila (Québec Artificial Intelligence Institute), University of Montreal, Montreal, Canada**

Alex Hernandez-Garcia

**Max Planck Institute for Intelligent Systems, Tübingen, Germany**

Stefan Bauer

**Gut Microbes & Health, Quadram Institute Bioscience, Norwich Research Park, Norwich, Norfolk, UK.**

Falk Hildebrand, Tamas Korcsmaros

**Earlham Institute, Norwich Research Park, Norwich, Norfolk, UK.**

Falk Hildebrand, Tamas Korcsmaros

**Centre Hospitalier Universitaire Sainte-Justine, Montréal, Canada.**

Prévost Jantchou

## Supplemental Tables

**Supplemental Table 1.** Model performance on sample and patient demographics. Samples were grouped by the five different metadata categories and the classification performance with the indicated metric determined for control and CD or UC (depending on the metadata group). Coefficient refers to the corresponding independent variable’s coefficient for the logistic regression function and SE refers to the standard error of the coefficient. **** indicates p-value < 0.0001, *** indicates p-value < 0.001, ** indicates p-value < 0.01, and * indicates p-value < 0.05.

| Metric | Group | Variable | Coefficient | SE |
| --- | --- | --- | --- | --- |
| F1 Score | Sample Type | Biopsy (vs. Stool) | -0.33 | 0.21 |
| F1 Score | Life Stage | Adult (vs. Pediatric) | 0.47 ** | 0.18 |
| F1 Score | BMI Stratification | BMI <30 (vs. BMI > 30) | 0.49 ** | 0.19 |
| F1 Score | Sex | Female (vs. Male) | 0.31 | 0.19 |
| F1 Score | IBD Type | CD (vs. UC) | 0.54 ** | 0.18 |
| AUC | Sample Type | Biopsy (vs. Stool) | -0.18 | 0.2 |
| AUC | Life Stage | Adult (vs. Pediatric) | 1.25 **** | 0.17 |
| AUC | BMI Stratification | BMI <30 (vs. BMI > 30) | -0.17 | 0.18 |
| AUC | Sex | Female (vs. Male) | 0.1 | 0.18 |
| AUC | IBD Type | CD (vs. UC) | 0.07 | 0.17 |
| Accuracy | Sample Type | Biopsy (vs. Stool) | -0.7 *** | 0.2 |
| Accuracy | Life Stage | Adult (vs. Pediatric) | 1.03 **** | 0.18 |
| Accuracy | BMI Stratification | BMI <30 (vs. BMI > 30) | 0.41 * | 0.19 |
| Accuracy | Sex | Female (vs. Male) | 0.2 | 0.19 |
| Accuracy | IBD Type | CD (vs. UC) | 0.17 | 0.18 |

**Supplemental Table 2.** Overview of QIIME2 processing for 15 microbiome datasets. Samples were collected from the listed ENA accession, with only samples corresponding to individuals in North America retained. Trim length was used as input for the trunc_len parameter, forward trim as the trim_left input for single end read and trim_left_f for paired-end reads, and reverse trim as the trim_left_r input for paired-end reads in the python API for QIIME2’s Dada2 plugin.

| Study ID | ENA Accession | Hypervariable Region | Trim Length | Forward Trim | Reverse Trim |
| --- | --- | --- | --- | --- | --- |
| American Gut | ERP012803 | V4 | 124 | 0 | 0 |
| CVDF | PRJNA308319 | V3-V4 | 290 | 40 | 40 |
| GEVERSC | PRJEB13680 | V4 | 174 | 0 | 0 |
| GEVERSM | PRJEB13679 | V4 | 174 | 0 |  |
| GLS | PRJEB23009 | V4 | 99 | 0 |  |
| HMP | ibdmdb.org | V4 | 249 | 0 | 0 |
| MUC | PRJNA317429 | V4 | 174 | 19 | 21 |
| PRJNA418765 | PRJNA418765 | V4 | 245 | 0 | 3 |
| PRJNA436359 | PRJNA436359 | V4 | 170 | 0 | 3 |
| QIITA10184 | PRJEB13895 | V4 | 120 | 0 |  |
| QIITA10342 | PRJEB13619 | V4 | 100 | 0 |  |
| QIITA10567 | PRJEB14674 | V4 | 99 | 0 |  |
| QIITA1448 | PRJEB13051 | V4 | 99 | 0 |  |
| QIITA2202 | PRJEB6518 | V4 | 99 | 0 |  |
| QIITA550 | PRJEB19825 | V4 | 149 | 0 |  |

## Supplemental Figures

**Supplemental Figure 1.**
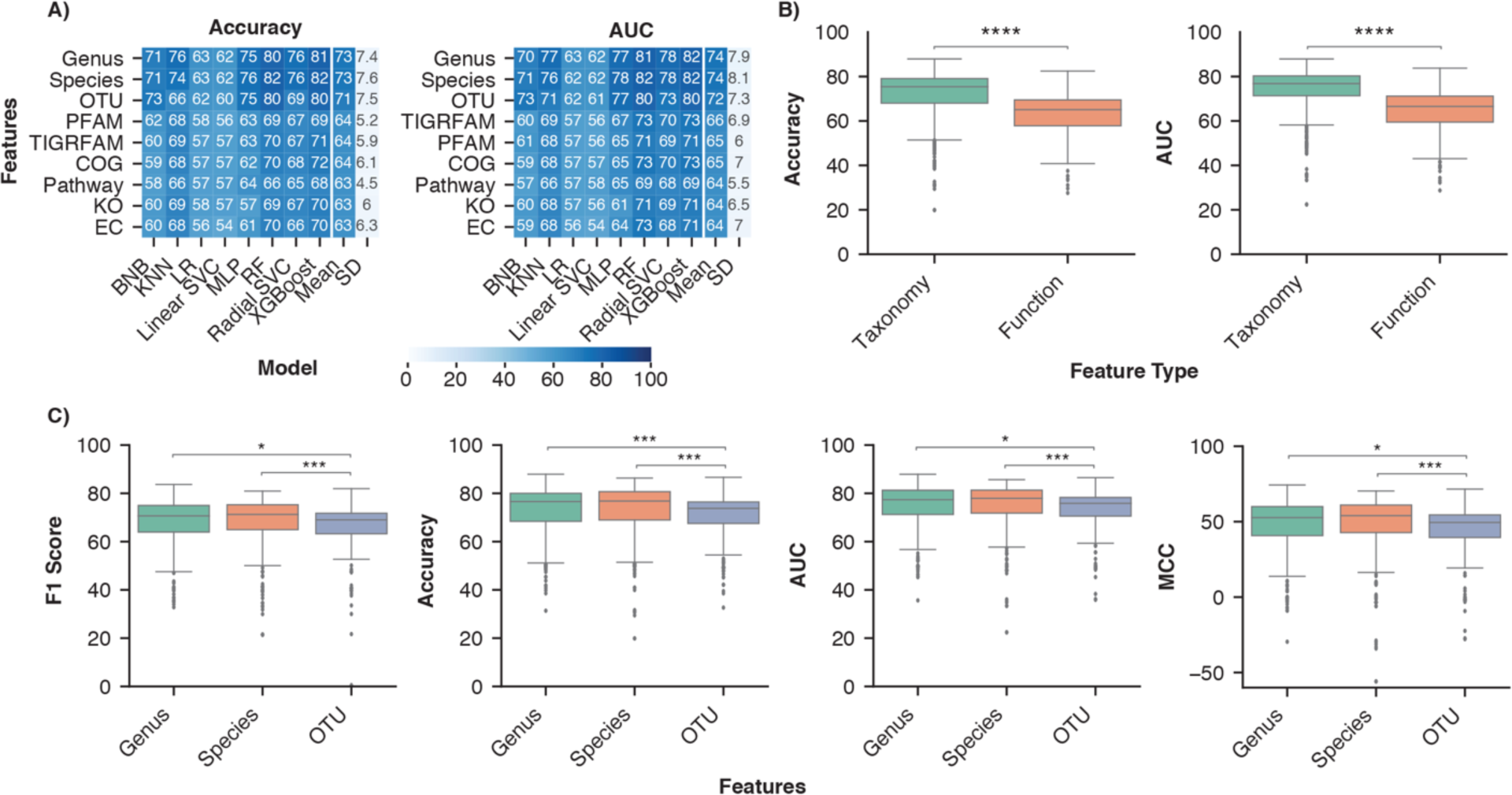
Greater classification of IBD samples with taxonomic features than functional features. A) Average performance for each type of feature across different model architectures. Rows were sorted in descending order by the mean column followed by the standard deviation (SD) column. B) Distribution of performance metrics across all normalization, batch effect reduction, and model combinations. Independent Mann-Whitney U-tests were performed to compare aggregate performance measured by each metric of taxonomy and functional features. C) Comparison of classification performance with the three taxonomic feature sets. All pairwise comparisons were performed with a Mann-Whitney U-test followed by Bonferroni correction and the significant comparisons are indicated. The analysis was limited to normalization (ILR, CLR, VST, ARS, LOG, TSS, NOT) and batch effect reduction (No batch reduction or Zero-Centering) methods that were performed on all feature sets. **** indicates p-value < 0.0001, *** indicates p-value < 0.001, * indicates p-value < 0.05.

**Supplemental Figure 2.**
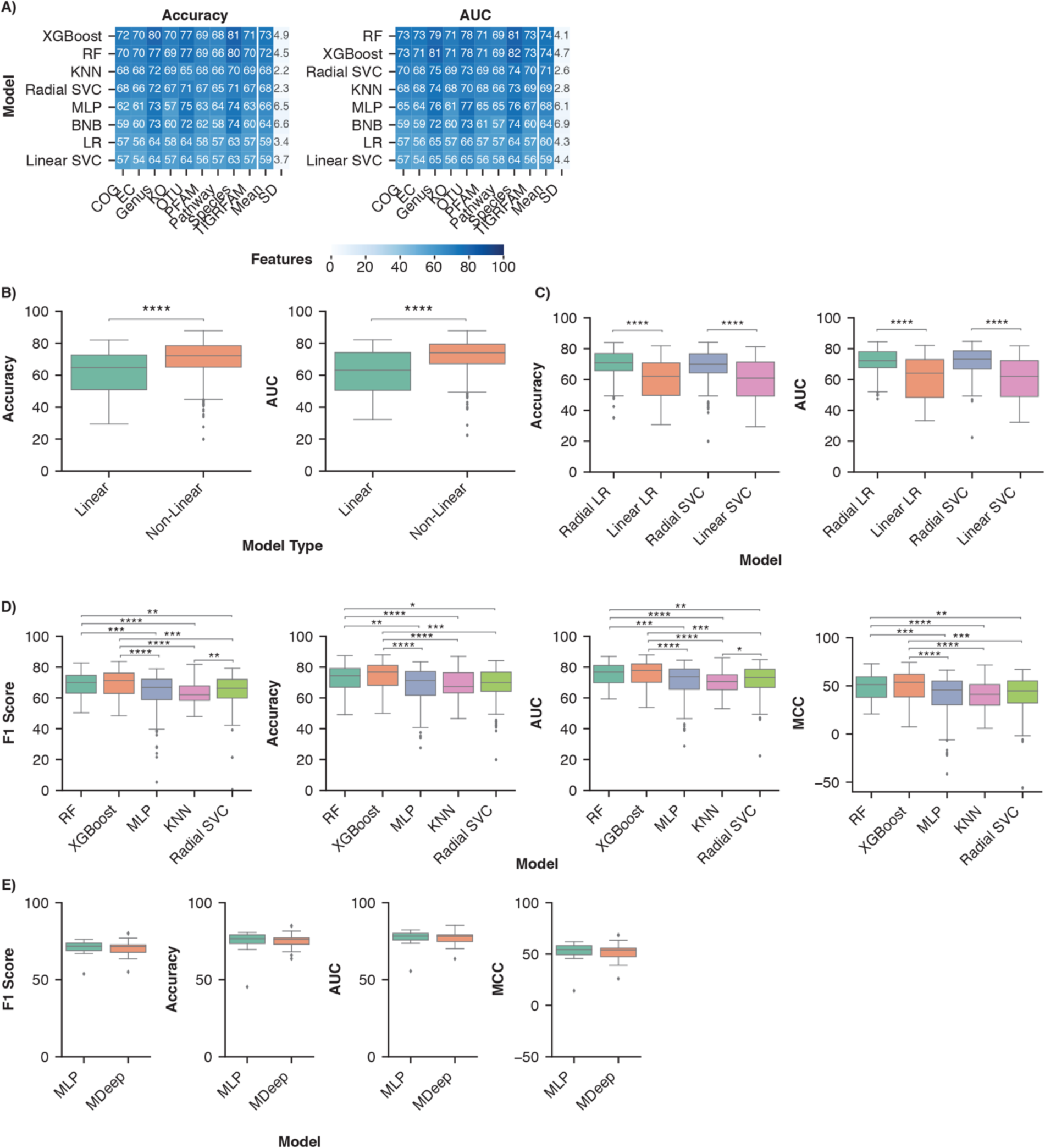
Greater IBD classification performance with non-linear than linear ML models. A) Average model performance for each feature set across normalization and batch effect reduction methods. B) Distribution of performance of non-linear and linear models. Independent Mann-Whitney U-tests were performed to compare each performance metric. Analysis was limited to datasets preprocessed using normalization (ILR, CLR, VST, ARS, LOG, TSS, NOT) and batch effect reduction (No batch reduction or Zero-Centering) methods performed on all feature types. C) Distribution of classification performance with the non-linear and linear variations of logistic regression and support vector machines across all feature sets. A Mann-Whitney U test with Bonferroni correction was performed to compare the linear and non-linear variation of each model respectively. D) Comparison of IBD classification performance between the non-linear models. All pairwise comparisons were performed by Mann-Whitney U test with a Bonferroni correction and the significant comparisons were labelled. E) Comparison of two neural network architectures: the convolutional neural network MDeep or a MLP. A Mann-Whitney U test was used to compare each performance metric and the significant comparisons were labelled. **** indicates p-value < 0.0001, *** indicates p-value < 0.001, ** indicates p-value < 0.01, and * indicates p-value < 0.05.

**Supplemental Figure 3.**
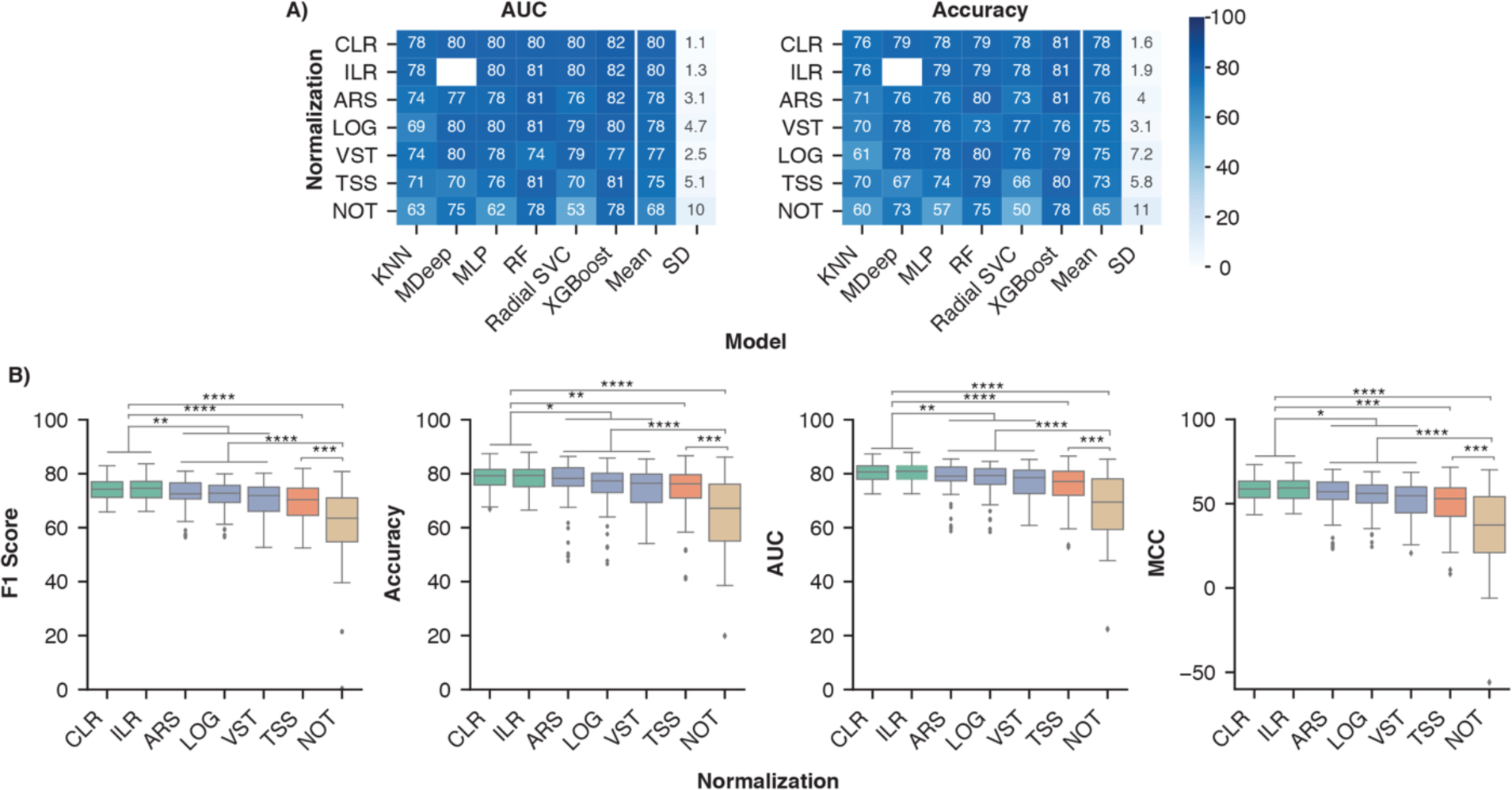
Compositional normalization methods lead to the highest model performance for IBD classification. A) Average model performance with each normalization method across all batch effect reduction methods. B) Comparing the effect of different classes of normalization methods. The compositional category consists of CLR and ILR (green), variance/distribution modifiers consists of VST, ARS, and LOG (blue), scaling consists of TSS (orange), and no normalization consists of NOT (brown). All pairwise combinations were compared with a Mann-Whitney U test with a Bonferroni correction and the significant comparisons labelled. **** indicates p-value < 0.0001, *** indicates p-value < 0.001, ** indicates p-value < 0.01, and * indicates p-value < 0.05.

**Supplemental Figure 4.**
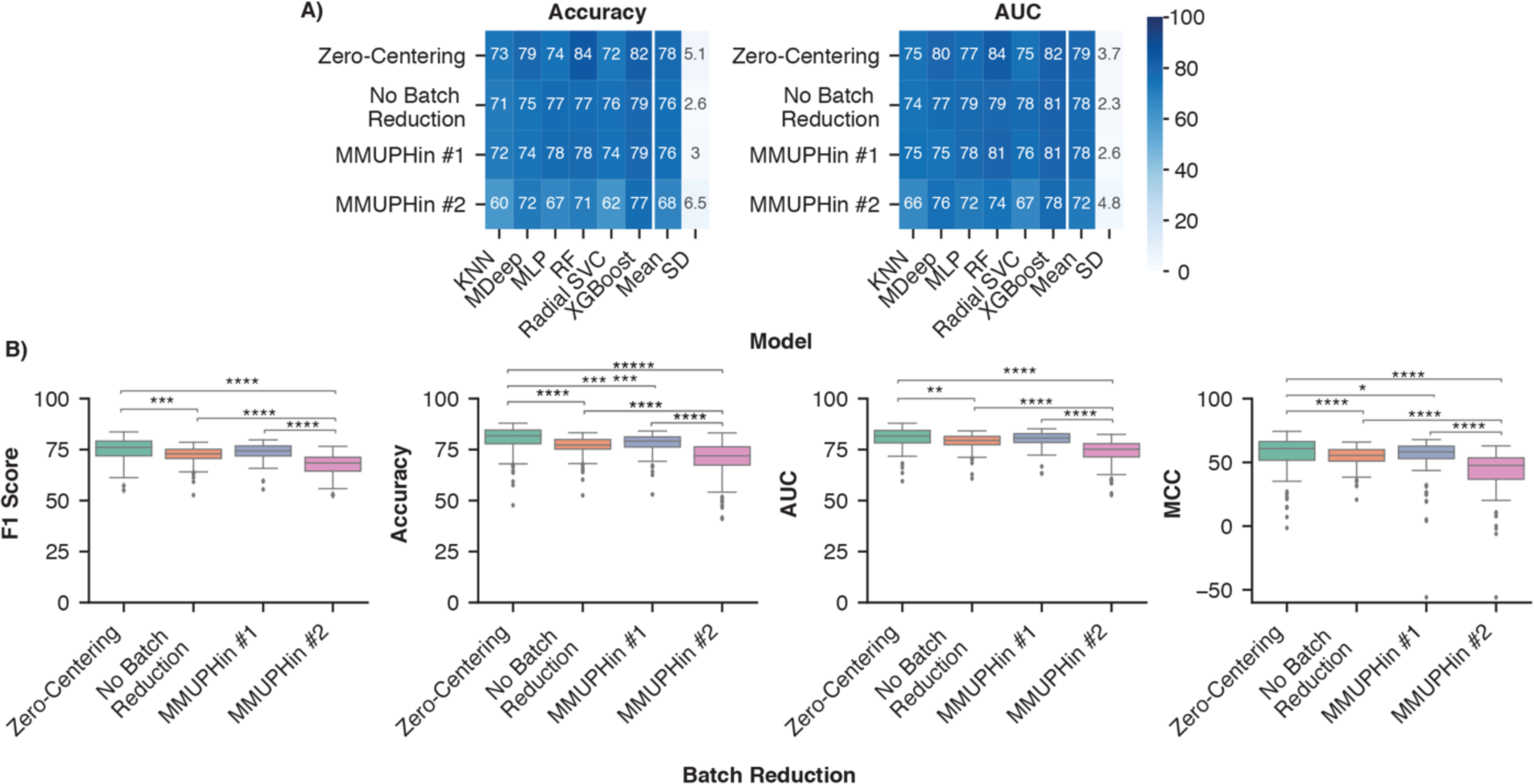
Removing batch effects with zero-centering improved IBD classification. A) Batch effect reduction methods sorted by average performance across all combinations normalization methods, taxonomic features, and non-linear ML models. B) Comparing the effect of different batch effect reduction methods on classification of IBD samples. All pairwise combinations were compared with a Mann-Whitney U test and the significant comparisons were labelled. **** indicates p-value < 0.0001, *** indicates p-value < 0.001, ** indicates p-value < 0.01, and * indicates p-value < 0.05.

**Supplemental Figure 5.**
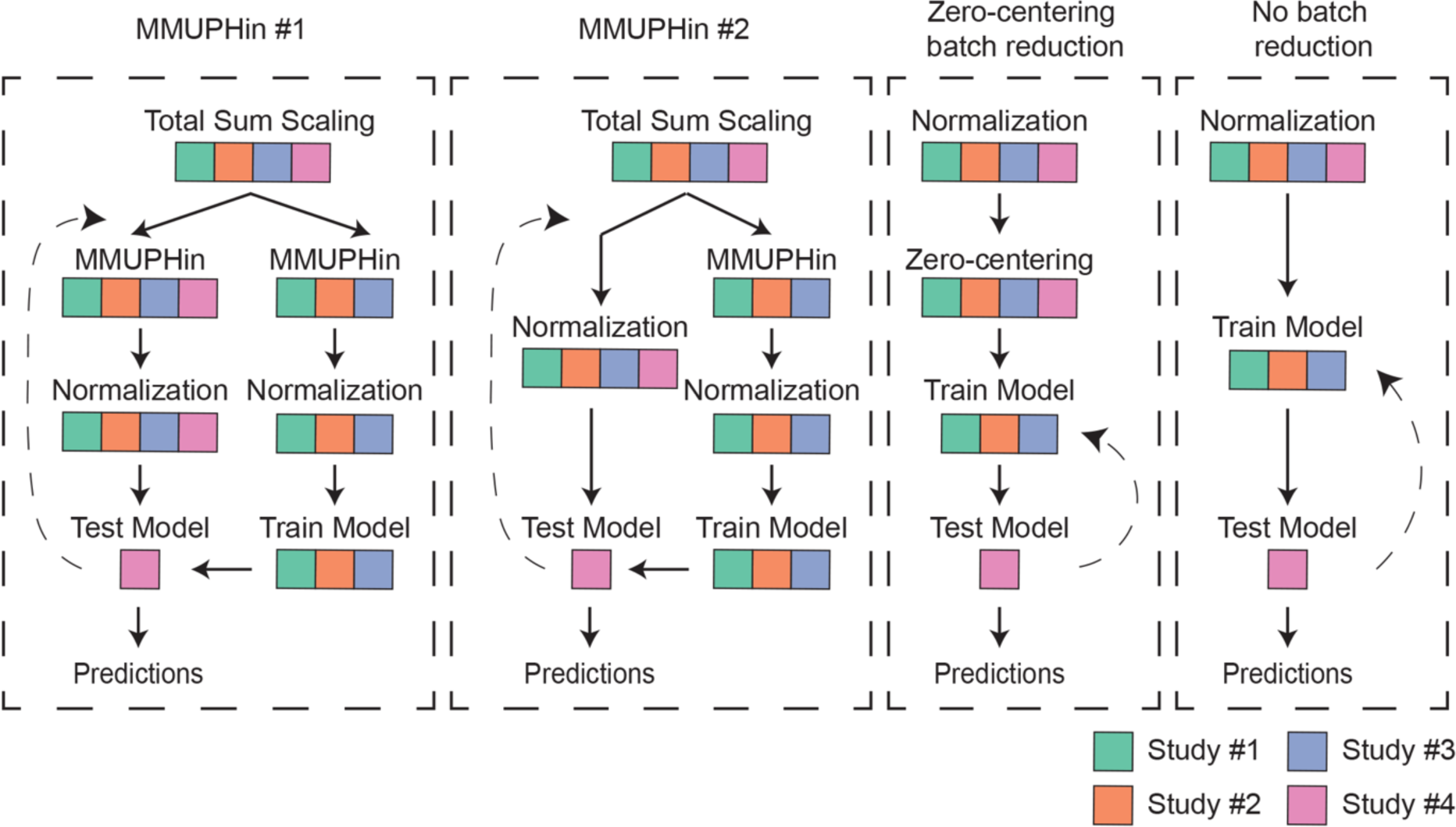
LODO cross validation pipeline for different batch effect reduction methods. Pipeline performance (combinations of a normalization method, batch effect reduction method, and ML model) was determined with LODO cross validation. Four variations were used to account for different requirements in the batch effect reduction step to ensure the training and test set were independent. The coloured boxes under the text indicate which studies the corresponding step was performed with. The diagram illustrates how a single dataset (illustrated by the different colours) is removed from the dataset the model is trained with and then tested on. The dashed arrow indicates the step returned to in each iteration, with a different dataset removed in each iteration.

